# Transcriptomic changes resulting from *STK32B* overexpression identifies pathways potentially relevant to essential tremor

**DOI:** 10.1101/552901

**Authors:** Calwing Liao, Faezeh Sarayloo, Veikko Vuokila, Daniel Rochefort, Fulya Akçimen, Simone Diamond, Alexandre D. Laporte, Dan Spiegelman, Qin He, Hélène Catoire, Patrick A. Dion, Guy A. Rouleau

**Author notes:** **Corresponding Author**: Dr. Guy A. Rouleau, MD, PhD, FRCPC, OQ, Department of Neurology and Neurosurgery, McGill University, Montréal, Québec, Canada H3A 2B4.

## Abstract

Essential tremor (ET) is a common movement disorder that has a high heritability. A number of genetic studies have associated different genes and loci with ET, but few have investigated the biology of any of these genes. *STK32B* was significantly associated with ET in a large GWAS study and was found to be overexpressed in ET cerebellar tissue. Here, we overexpressed *STK32B* in human cerebellar DAOY cells and used an RNA-Seq approach to identify differentially expressed genes by comparing the transcriptome profile of these cells to the one of control DAOY cells. Pathway and gene ontology enrichment identified axon guidance, olfactory signalling and calcium-voltage channels as significant. Additionally, we show that overexpressing *STK32B* affects transcript levels of previously implicated ET genes such as *FUS.* Our results investigate the effects of overexpressed *STK32B* and suggest that it may be involved in relevant ET pathways and genes.

## Introduction

Essential tremor (ET) is one of the most common movement disorders and is typically characterized by a kinetic tremor in the hand or arms^1^. The severity tends to increase with age and may involve different regions such as the head, voice or jaw^2^. Previous studies investigating the histology of post-mortem tissue have pinpointed the cerebellum as a region of interest for ET^3^. Specifically, abnormalities with Purkinje cell axons and dendrites were found in ET brains^4^. Furthermore, the olivocerebellar circuitry has been implicated in ET pathology and many calcium voltage channels are highly expressed in this circuitry^5^.

The genetic etiology of ET has remained largely elusive and most studies have focused on common or rare variants and on looking for genetic overlap with other disorders^6–7^. Twin studies have shown that ET has a concordance of 69–93% in monozygotic twins and 27–29% in dizygotic twins, which suggests that both genetic and environmental factors drive the onset and development of this complex trait^8^. A recent genome-wide association study (GWAS) identified a significant locus in *STK32B* and found that ET patients overexpressed *STK32B* in cerebellar tissue by comparison to healthy controls^9^. *STK32B* is transcribed and translated into YANK2, a serine/threonine kinase, that has not been well characterized. There have been several exome-wide studies that implicated different genetic variants as causes of ET. The first ET-implicated gene found through exome sequencing was the Fused in Sarcoma gene (FUS)^10^. However, it is unclear whether these “ET” genes interact or have indirect effects on the expression of each other.

To understand the effects of overexpressed *STK32B* and identify pathways potentially relevant to ET, we overexpressed the gene in human cerebellar DAOY cells and compared transcriptomic changes in overexpressed cells and empty-vector controls using RNA sequencing. Several interesting pathways such as axon guidance, calcium ion transmembrane transport, and olfactory transduction were significantly enriched after overexpression of *STK32B.* We also identified previously implicated ET genes whose expression is dysregulated through the overexpression of *STK32B,* suggesting that overexpressed *STK32B* may have relevant downstream effects.

## Results

### Confirming overexpression of *STK32B*

The *STK32B* stable cell lines had higher RNA expression of *STK32B,* which was detected by reverse-transcriptase qPCR (Supplementary Figure 1). RNA sequencing (RNA-Seq) also confirmed that the stable cell lines have higher *STK32B* RNA levels compared to controls; it was the most significantly differentially expressed gene (Figure 1–2).

**Figure 1.**
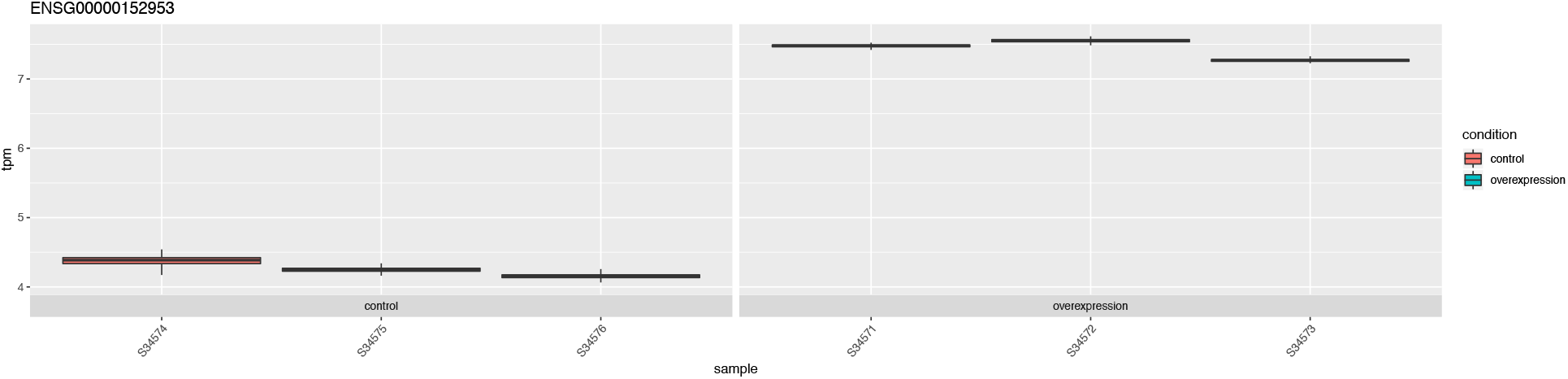
The RNA expression of *STK32B* overexpressed cells compared to empty-vector controls based on RNA sequencing data. Error bars show the technical variance determined with 200 bootstraps. Abbreviations: TPM, transcript per million.

**Figure 2.**
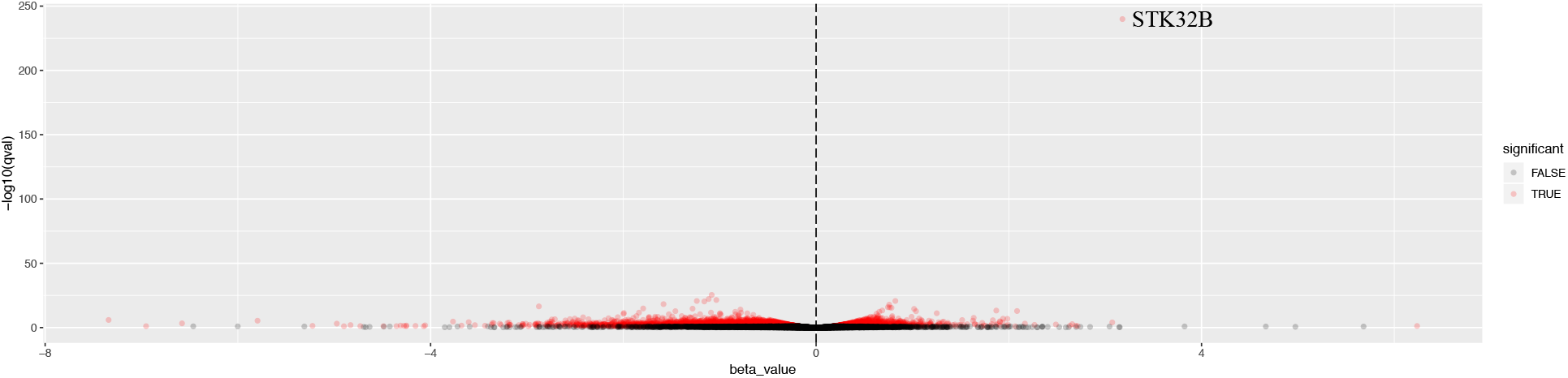
Volcano plot of comparing overexpressed *STK32B* to empty-vector controls. Differentially expressed genes (q < 0.05) are shown in red. Beta value was determined with a Wald’s test.

### Quality control of RNA-seq analyses

The QQ-plot for differentially expressed genes did not show stratification or potential biases (Supplementary Figure 2). The M-A plot showed slight enrichment of downregulated genes compared to upregulated genes (Supplementary Figure 3). A principal component (PC) plot showed separation of controls and *STK32B* cell lines (Figure 3). Additionally, the dendrogram with heatmap shows that controls and *STK32B* cell lines are distinct (Supplementary Figure 4). The mean-variance plot shows the shrinkage under the sleuth model (Supplementary Figure 5).

**Figure 3.**
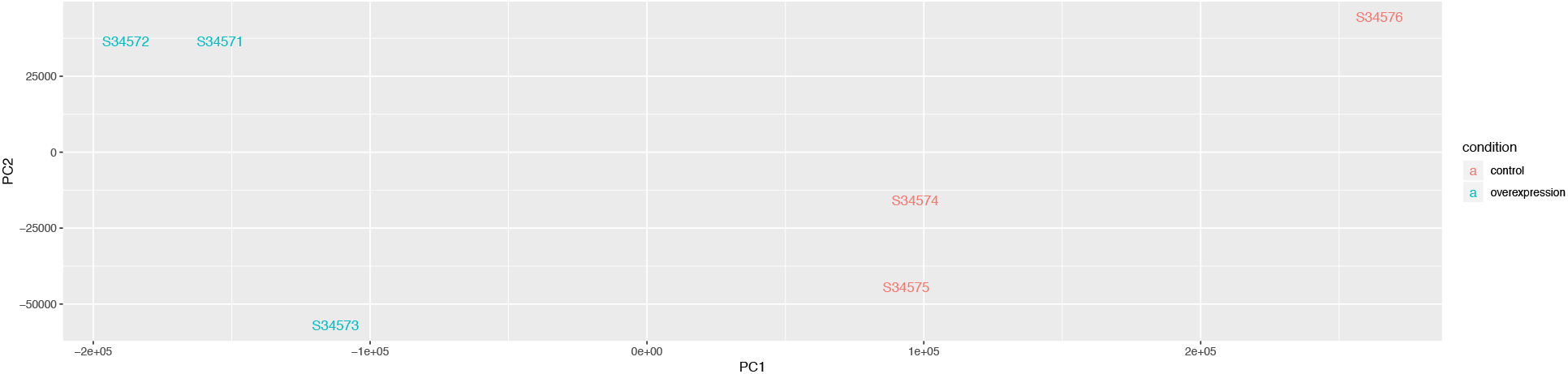
Principle component analysis plot of RNA sequencing data of *STK32B* overexpressed cells and empty-vector controls. Overexpressed cells shown in blue and controls shown in red.

### Overexpression of *STK32B* affects expression of previously iden;fied ET-associated genes

A total of 3,794 genes were found to be differentially expressed with a q-value <0.05. There were 425 genes with a β > |0.5| and q-value <0.05. Amongst the 425 genes, three potentially relevant ET genes were dysregulated: *FUS* (q-value = 0.007, β=0.812) and two calcium voltage channel genes that are enriched in the olivocerebellar circuitry, *CACNA1C* (q-value = 0.011, β= 0.503) and *CACNA1A* (q-value = 0.030, β= 0.702). Several pathways and gene ontologies were enriched for differentially expressed genes (DEGs) due to the overexpression of *STK32B*, including olfactory transduction (P=2.k8E-39), axon guidance (P=9.50E-36), and calcium ion transmembrane transport (P=3.02E-31) (Table 1).

**Table 1.** Gene ontology and pathway analyses of differentially expressed genes.

| Gene Set ID | Gene Set Name | *P* | Database |
| --- | --- | --- | --- |
| KEGG_OLFACTORY_TRANSDUCTION | Olfactory transduction | 2.68E-39 | KEGG |
| KEGG_PATHWAYS_IN_CANCER | Pathways in cancer | 2.96E-28 |  |
| KEGG_FOCAL_ADHESION | Focal adhesion | 1.19E-26 |  |
| KEGG_BLADDER_CANCER | Bladder cancer | 6.53E-25 |  |
| KEGG_ECM_RECEPTOR_INTERACTION | Ecm-receptor interaction | 8.38E-25 |  |
| GO:0001650 | fibrillar center | 1.13E-29 | GO<br>Cellular |
| GO:0031012 | extracellular matrix | 8.57E-27 |  |
| GO:0001725 | stress fiber | 1.24E-26 |  |
| GO:0005925 | focal adhesion | 2.82E-23 |  |
| GO:0015629 | actin cytoskeleton | 4.02E-22 |  |
| REACTOME:R-HSA-381753 | Olfactory Signaling Pathway | 3.22E-46 | Reactome |
| REACTOME:R-HSA-418555 | G alpha (s) signalling events | 1.19E-29 |  |
| REACTOME:R-HSA-3000170 | Syndecan interactions | 1.53E-27 |  |
| REACTOME:R-HSA-2022090 | Assembly of collagen fibrils and other multimeric structures | 4.31E-25 |  |
| REACTOME:R-HSA-1474290 | Collagen formation | 2.35E-24 |  |
| GO:0050911 | detection of chemical stimulus involved in sensory perception of smell | 1.00E-44 | Go<br>Processes |
| GO:0007411 | axon guidance | 9.50E-36 |  |
| GO:0048146 | positive regulation of fibroblast proliferation | 1.31E-33 |  |
| GO:0070588 | calcium ion transmembrane transport | 3.02E-31 |  |
| GO:0050907 | detection of chemical stimulus involved in sensory perception | 3.14E-31 |  |

**Table 2.**
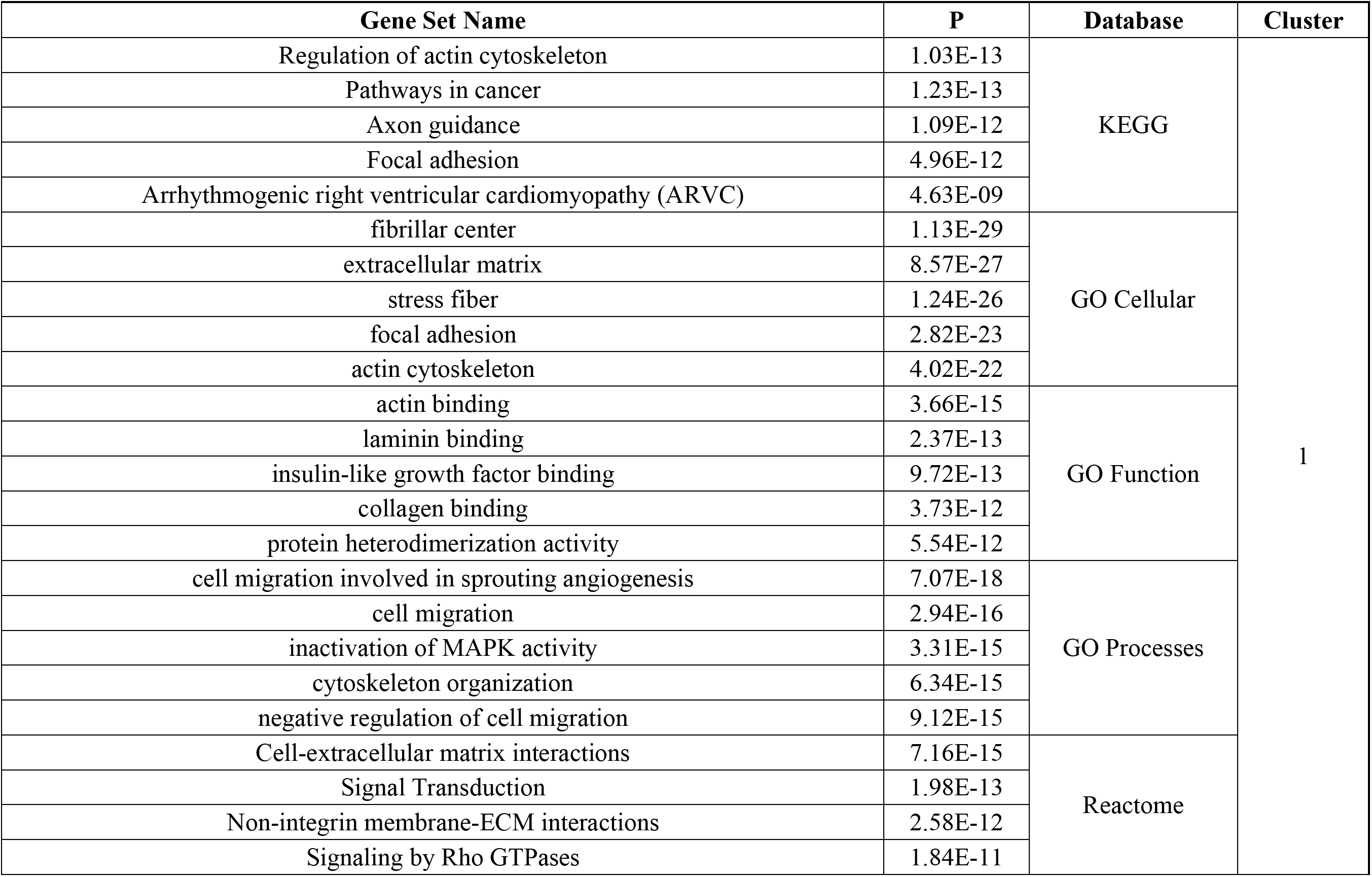
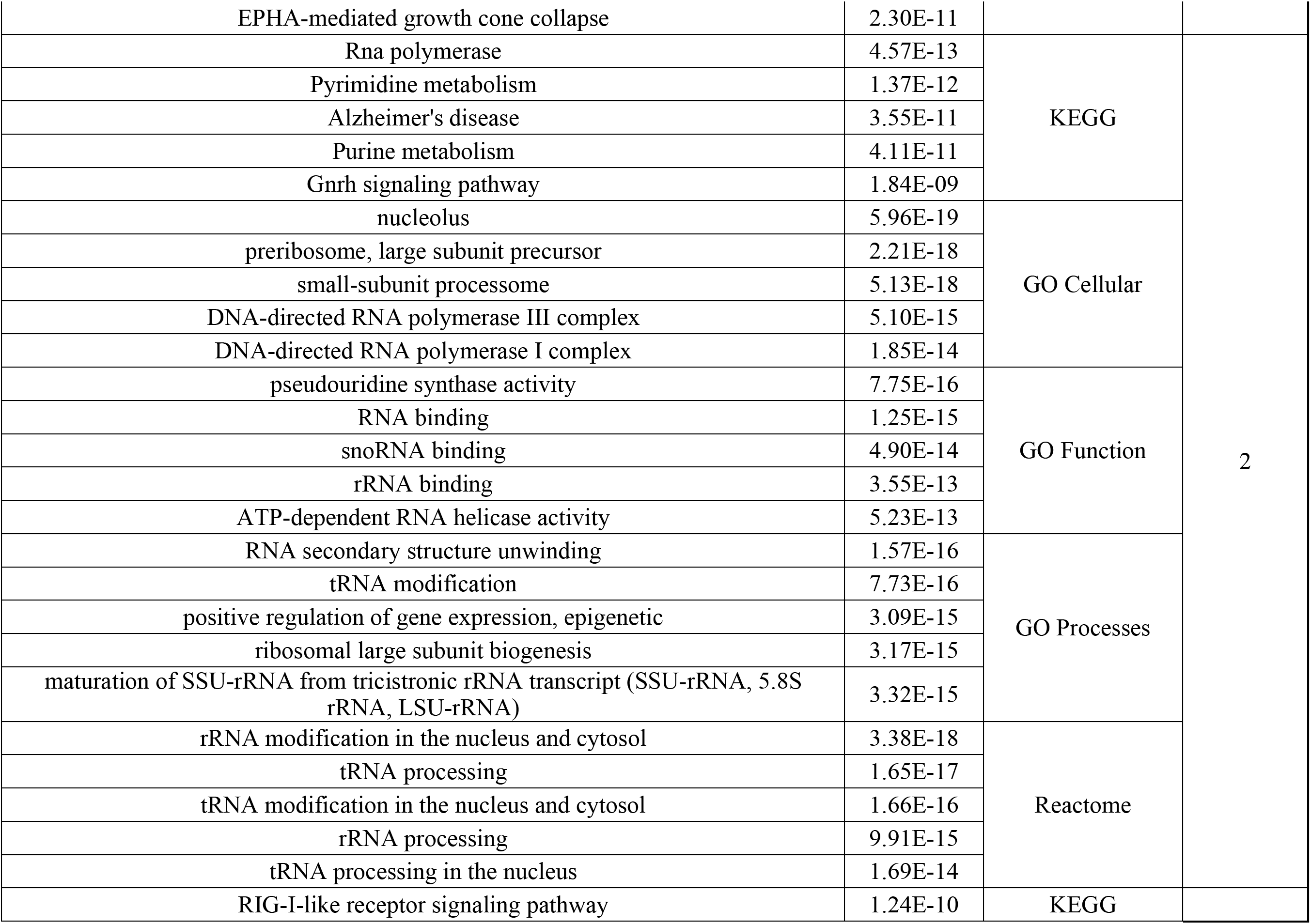
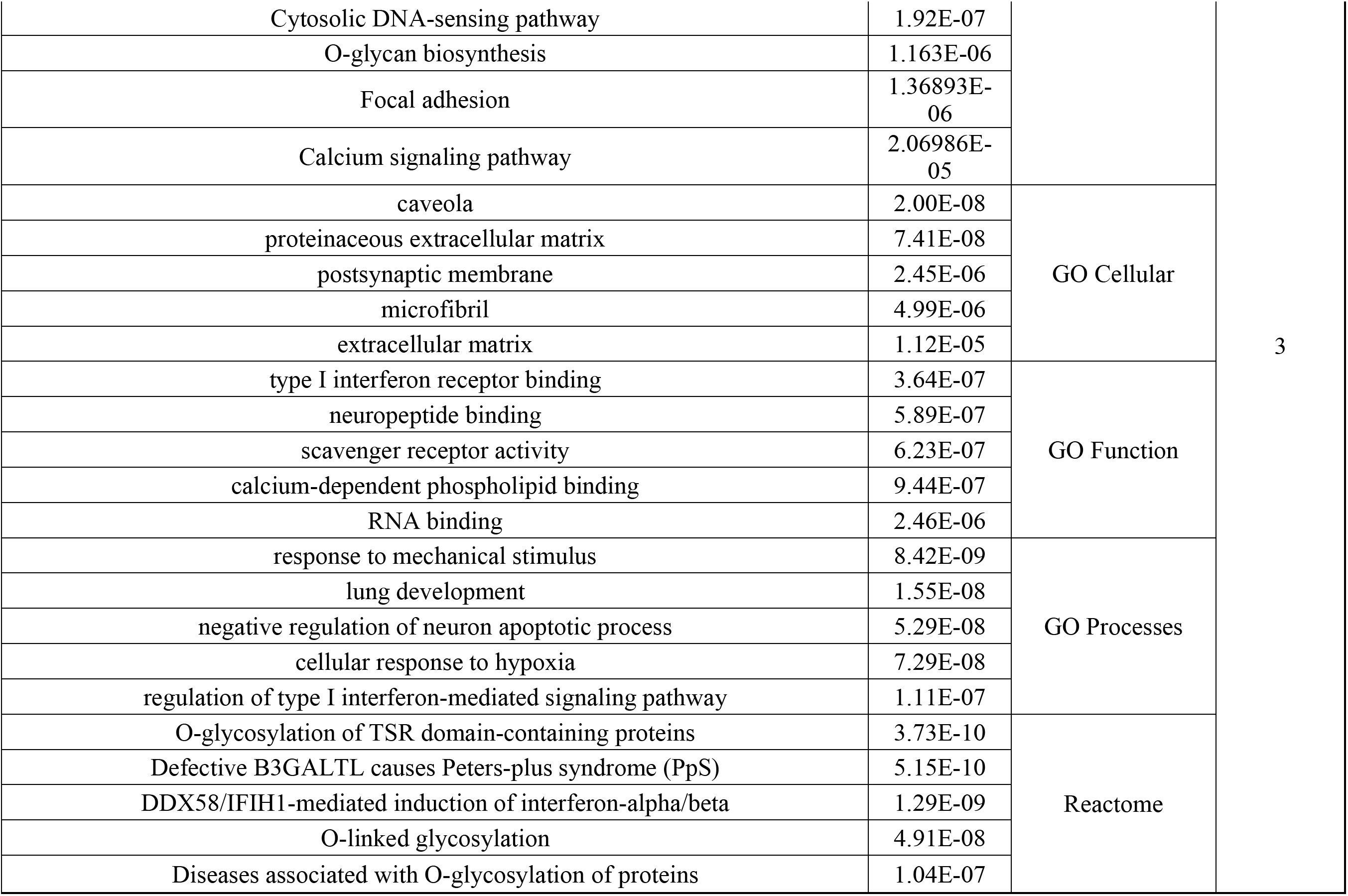
Gene ontology and pathway analyses of differentially expressed gene for the three largest clusters based on coexpression.

### Gene network analyses

To identify which cluster of DEGs drove the significant pathways, gene network analysis based on co-expression was done and identified 14 different clusters within the differentially expressed genes (Figure 4). Analysis of the top 3 largest clusters for pathway enrichments found several similar pathways in different databases including axon guidance (P<1.09E-12) in cluster 1 and calcium signalling pathway in cluster 3 (2.06E-05).

**Figure 4.**
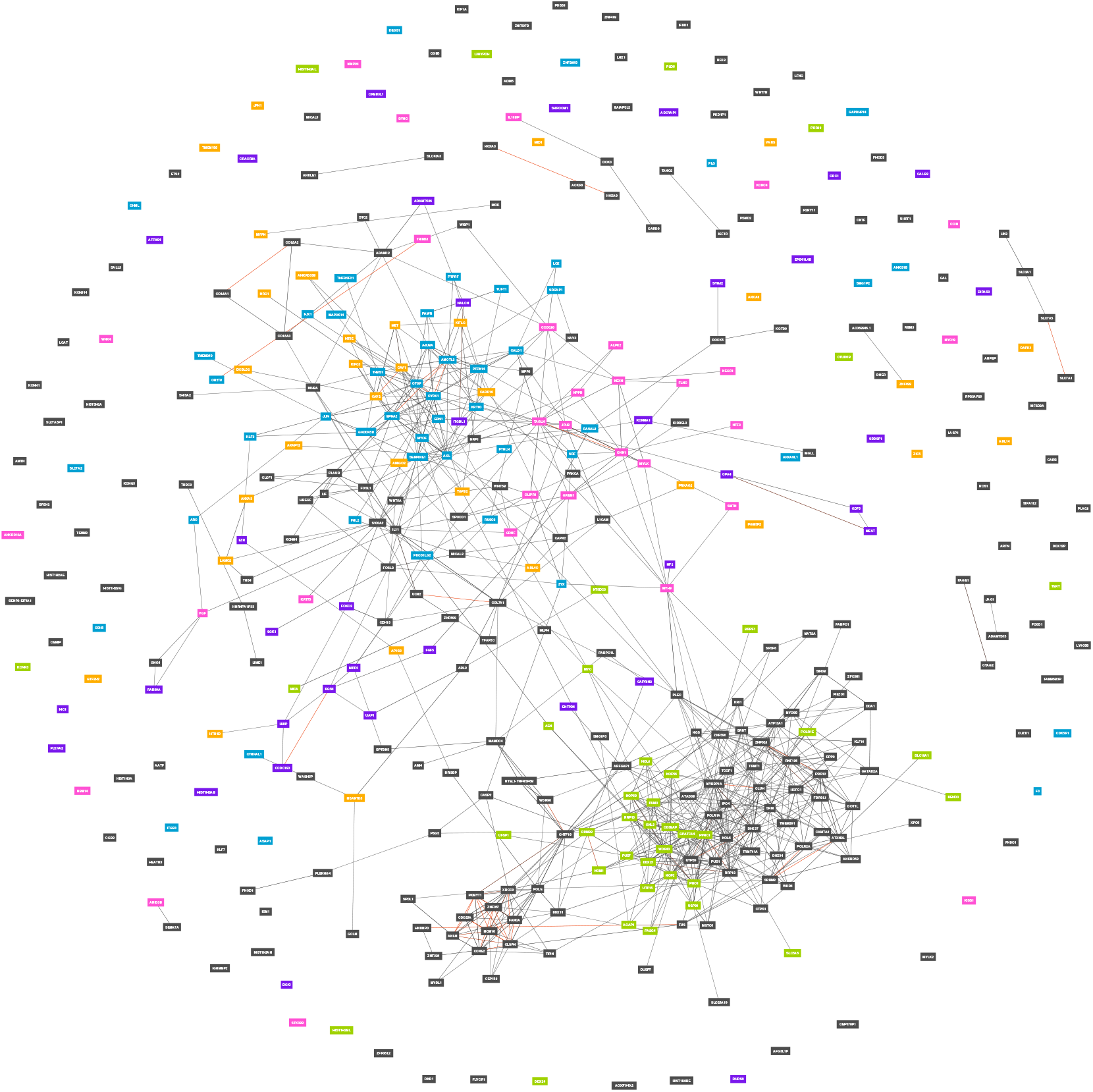
Gene network of differentially expressed genes based on co-expression of RNA sequencing data for 31,499 samples. Each unique colour represents a cluster.

## Discussion

The genetic etiology of ET is complex and likely explained by a combination of copy number variants, rare variants, gene-gene interactions and common variant drivers. One of the most significantly enriched pathways we identified was olfactory signalling and transduction. Previous reports have shown conflicting evidence about olfactory loss in ET^11,12^. It may be that ET patients with dysregulated olfactory signalling have overexpressed *STK32B*, which may contribute to the subset of ET patients with olfactory loss.

Interestingly, *FUS* was amongst the genes dysregulated when *STK32B* was overexpressed. In the exome familial ET study that identified *FUS,* we also found reduced mRNA levels for *FUS*^10^. Similarly, the expression of *FUS* is lower when *STK32B* is overexpressed, suggesting an indirect relationship between the two genes. Additionally, two calcium voltage-gated channel genes, *CACNA1C* AND *CACNA1A* were overexpressed. Enrichment analyses found calcium ion transmembrane transport to be highly significant in GO Processes, suggesting that *STK32B* may play a role upstream of these genes. In the olivocerebellar circuitry, a system implicated in ET, is enriched for both of these genes^5,13^. *CACNA1A* has been shown to be predominantly expressed in Purkinje cells, a cell type relevant to ET^13^. Another enriched pathway was axon guidance. A previous study identified the *TENM4* missense variant segregating within ET families^14^. *TENM4* is a regulator of axon guidance and myelination and TenmO knockout mice showed an ET phenotype, suggesting that axon guidance is important in ET^14^. Similarly, overexpressed *STK32B* could be dysregulating important pathways relevant to axon guidance.

By sub-stratifying the overexpressed genes into different clusters that are co-expressed, several interesting pathways involving the cardiovascular system were identified. Pathways such as right ventricular cardiomyopathy in KEGG and angiogenesis in GO Processes were found among the top five significant pathways. In certain ET patients, beta-blockers can reduce tremor magnitude and frequency. Beta-blockers lower blood pressure and are used to treat irregular heart rhythm and other cardiovascular phenotypes. A common treatment for ET is propranolol, a beta-blocker. This was a drug developed for cardiovascular health that affects adrenergic activity, but still reduces tremor in ET individuals. This could suggest that overexpressed *STK32B* affects cardiovascular health, which may in turn affect or lead to the ET phenotype or that *STK32B* may be pleiotropic and affect the nervous and cardiovascular system differently.

Due to the heterogeneity of ET, it is likely that not all ET-affected individuals have overexpression of *STK32B.* One limitation of this study is that DAOY cells are a cancerous cell line and may not be the ideal model for understanding the transcriptome of Purkinje cells. Future studies on the effects of *STK32B* overexpression could use induced pluripotent stem cells differentiated into cells with Purkinje cells properties; such cells would likely have less biological noise. However, our findings support the idea that *STK32B* is a gene of interest for a subset of ET cases, and further investigations into how *STK32B* may interact with other genes are warranted.

## Methods and Materials

### Cell Culture and plasmid construction

The DAOY cell line was cultured in Eagle’s Minimum Essential Medium (EMEM) with 10% fetal bovine serum (FBS) at 37 degrees Celsius and 5% CO2 with the Glico Pen-Strep-Glutamine cocktail at 1X. Cells were passaged every 2 days at 80-90% confluence and at the same time. The cDNA of *STK32B* (NM_001306082.1) was inserted into a pcDNA3.1(+) vector containing a neomycin resistance gene, used as a selectable marker. This transcript was picked because it had the highest expression in the cerebellum based on GTEx 53 v7. The plasmid was transformed and expanded in XL10-Gold *Escherichia coli* strain (Stratagene). The cDNA of the plasmid was sent for Sanger sequencing to validate the gene sequence.

### Transfection and deriving stable cell lines

The cells were transfected for 48 hours with 8 ug of DNA of using the jetPRIME transfection reagent (Polyplus). A green fluorescent protein (GFP) control vector was used to assess and estimate transfection efficiency and optimize transfection parameters. Stable cell lines were established by subjecting transfected cells to G418 antibiotic at 1 μg/mL for 10 days and maintained at a concentration of 400 μg/mL for an additional 2 weeks. A kill curve was done to determine the ideal antibiotic selection. In parallel, empty pcDNA3.1(+) vector control lines were also grown and underwent antibiotic selection.

### RNA extraction and sequencing

RNA was extracted using the RNeasy Mini Kit (Qiagen). The RNA concentration was measured using the Synergy H4 microplate reader. RNA for the top three overexpressed *STK32B* cell lines and three empty-vector controls was sent to Macrogen Inc. for sequencing. RT-qPCR was used to confirm overexpression in the cell liens compared to controls. Library preparation was done with the TruSeq Stranded Total RNA Kit (Ilumina) with Ribo-Zero depletion. Sequencing was done on the NovaSeq 6000 at 150bp paired-end reads with a total of 200M reads. Samples were randomized for cell lysis, RNA extraction, Ribo-Zero depletion, library preparation and sequencing to account for potential batch effects.

### Data processing and quality control

The FASTQ files were pseudo-aligned with Salmon using the Ensembl v94 annotation of the human genome^16^. The parameters of Salmon included the following: 200 bootstraps with mapping validation, GC bias correction, sequencing bias correction, 4 range factorization bins, a minimum score fraction of 0.95 and VB optimization with a VB prior of 1e-5. The estimated counts were analyzed using sleuth to identify DEGs^17^. A likelihood ratio test was used to identify differentially expressed transcripts and a Wald test was used to get β values (log_2_(x + 0.5)). Additionally, transcript-level aggregation was done against the Ensembl v94 annotation to determine gene-level differential expression. P-values were corrected using the Benjamini-Hochberg procedure to account for false discovery rate (FDR). Q-value (p-values corrected for FDR) significance was set for an FDR of <0.05. A principal component (PC) plot and heatmap was done to identify clusters.

### Pathway and gene set enrichment analysis

Pathway and gene set enrichment for relevant databases was done using Gene Network 2.0. To investigate genes with larger fold changes, only differentially expressed genes with a β > |0.5| were analyzed for pathway enrichments. Network clustering was done by grouping genes based on coexpression. Pathway analyses were also conducted with Gene Network 2.0 for the clusters using a Wilcoxin test.

## Acknowledgments

This work was supported by a Canadian Institutes of Health Research Foundation Scheme grant (#332971). G.A.R. holds a Canada Research Chair in Genetics of the Nervous System and the Wilder Penfield Chair in Neurosciences. C.L. is a recipient of the Frederick Banting and Charles Best Canada Graduate Scholarship from the Canadian Institutes of Health Research (CIHR).

## Data availability statement

Any data is available upon request of the corresponding author.

