## Supplementary material for "Transcriptomic changes resulting from *STK32B* overexpression identifies pathways potentially relevant to essential tremor": Suplementary File

**Supplementary Figures**

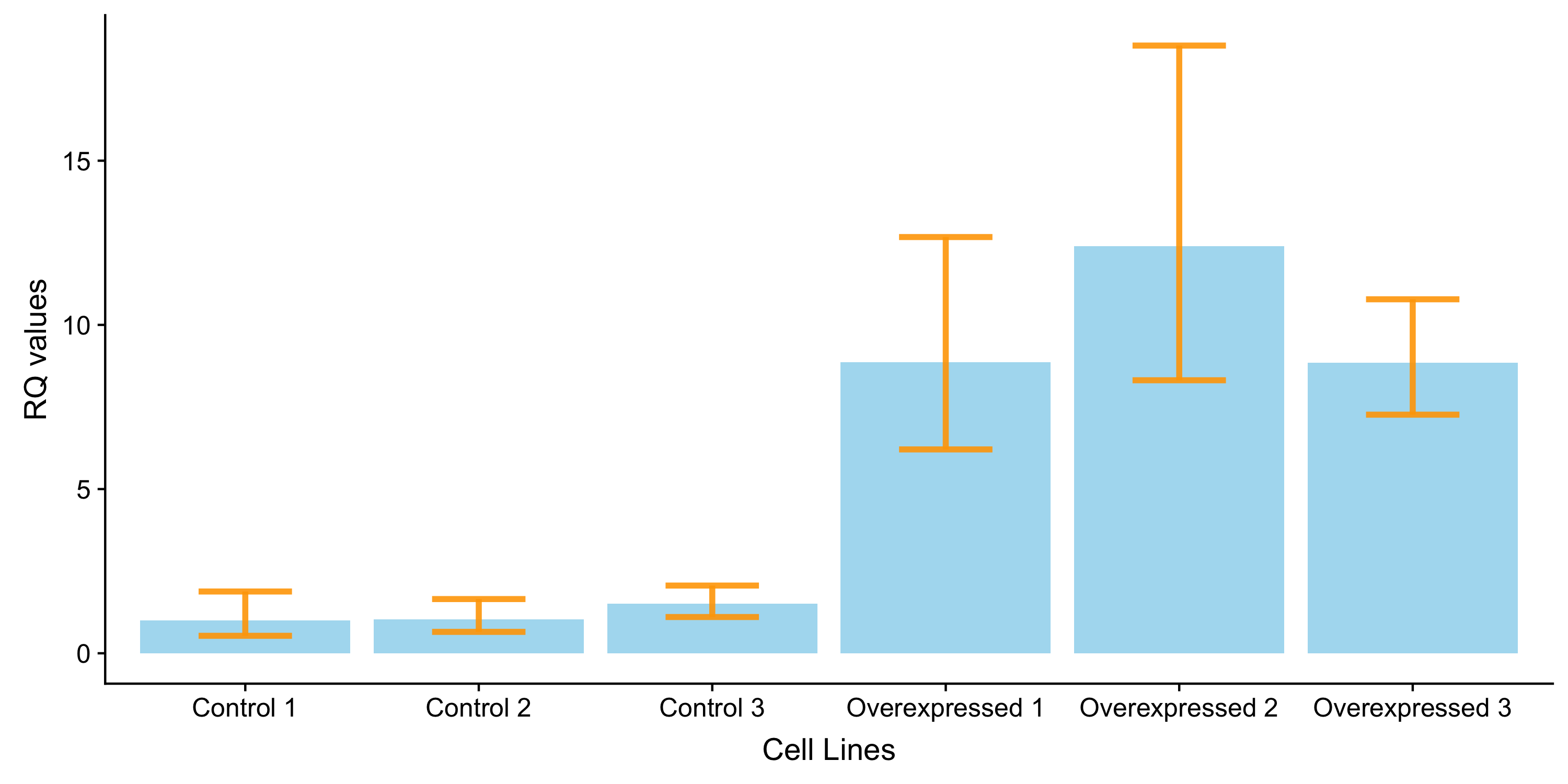

**Supplementary Figure 1. Relative quantification values of RNA sent for sequencing.** Samples were done in quadruplicate. Error bars represent min and max amongst the quadruplicates.

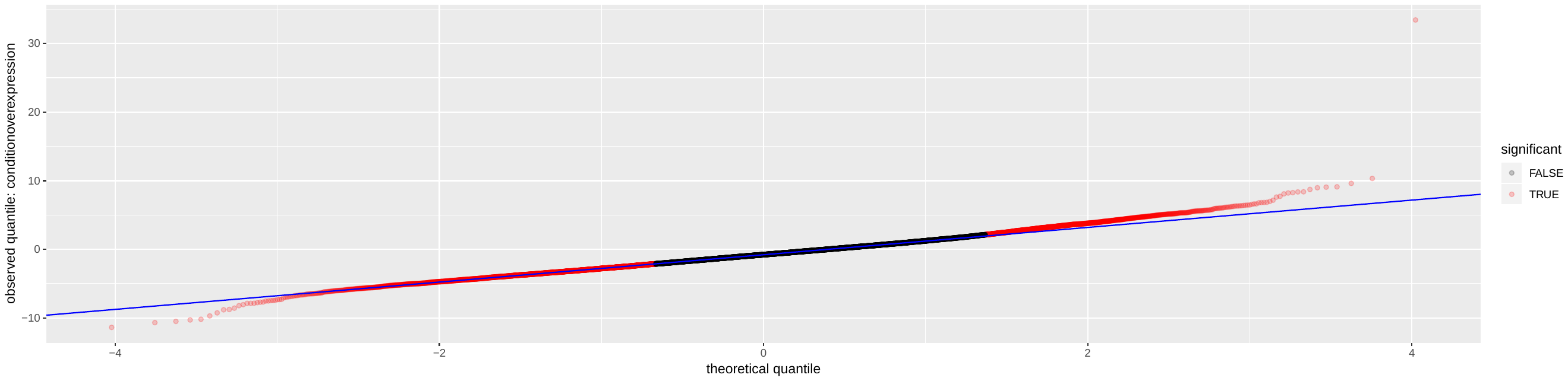

**Supplementary Figure 2. QQ-plot of differentially expressed genes for RNA sequencing data.**

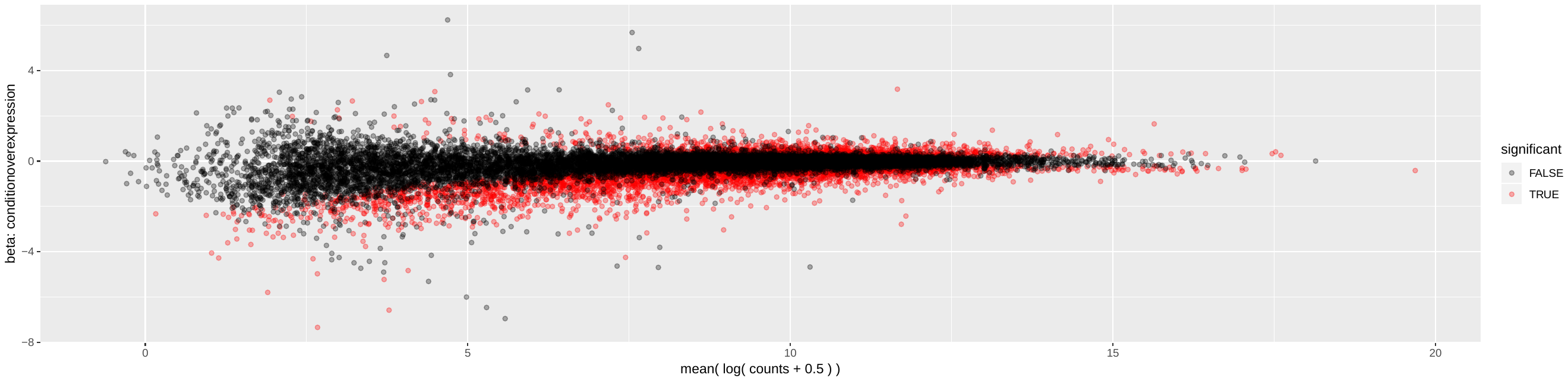

**Supplementary Figure 3. MA-plot of RNA sequencing data.**

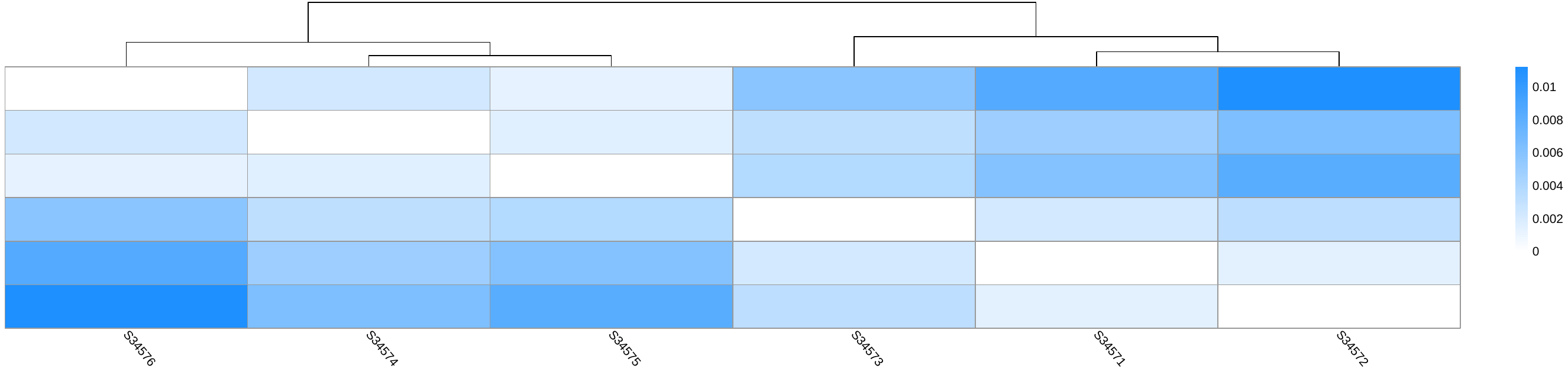

**Supplementary Figure 4. Heatmap of RNA sequencing data for both controls and overexpressed.** S34571, S34572 and S34573 were overexpressed cells. S34574, S34575 and S34576 were controls.

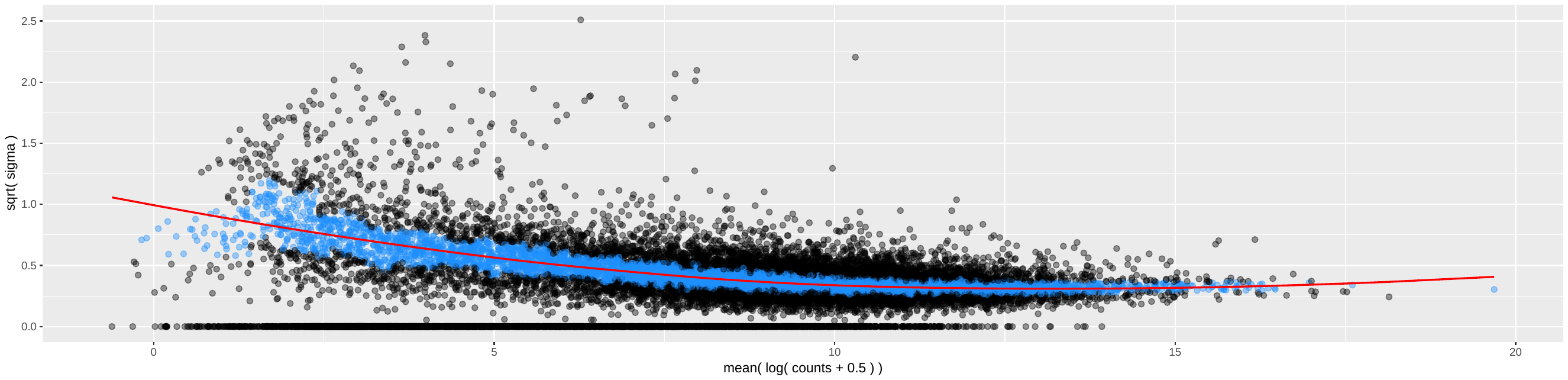

**Supplementary Figure 5. Mean-variance plot of the differentially expressed data.** Data was processed through sleuth for differential expression.
